# Oral Administration of the carboxylic acid ester prodrug NHC SN_9 protects mice from lethal Orthopoxvirus challenge: Implications for the Treatment of Monkeypox

**DOI:** 10.64898/2025.12.30.696971

**Authors:** Andrei E. Siniavin, Vladimir A. Gushchin, Natal’ya S. Shastina, Elizaveta S. Darnotuk, Sergey I. Luyksaar, Inna Dolzhikova, Anna M. Inshakova, Nailya A. Zigangirova, Alexander L. Gintsburg, Denis Y. Logunov

**Affiliations:** Federal State Budget Institution “National Research Centre for Epidemiology and Microbiology Named after Honorary Academician N. F. Gamaleya” of the Ministry of Health of the Russian Federation, Moscow, Russia; Department of Medical Genetics, I. M. Sechenov First Moscow State Medical University, Moscow, Russia; Institute of Fine Chemical Technologies, MIREA—Russian Technological University, 119571, Moscow, Russia

## Abstract

The global resurgence of monkeypox (Mpox) since 2022 has highlighted an urgent need for effective antiviral therapeutics against orthopoxvirus infections. While vaccines provide prophylactic protection, therapeutic options remain limited, particularly for immunocompromised individuals. Previously, we demonstrated potent antiviral activity of N4-hydroxycytidine (NHC) analogs against coronaviruses, including SARS-CoV-2. In the present study, we evaluated the antiviral efficacy of one such analog, NHC conjugate with phenylpropionic acid – SN_9, against orthopoxviruses using Vaccinia virus (VACV) as a model. SN_9 demonstrated superior *in vitro* antiviral activity compared with EIDD-2801 (molnupiravir), significantly inhibiting VACV replication. *In vivo*, oral administration of SN_9 provided substantial protection in murine models of orthopoxvirus infection, achieving 90% survival in a sublethal infection model and 70% survival in a lethal challenge model. Animals receiving treatment showed a reduction in disease severity and faster weight recovery. Given the high genetic and antigenic similarity among orthopoxviruses, including VACV, variola virus, and monkeypox virus (MPXV), these findings suggest that SN_9 represents a promising candidate for the treatment and prophylaxis of Mpox.

## Introduction

Monkeypox virus (MPXV), a member of the Orthopoxvirus genus, has emerged as a significant global public health concern following the unprecedented multi-country outbreak that began in 2022. Over 110 countries reported thousands of confirmed cases and hundreds of fatalities, marking the fifth recorded outbreak of Mpox and underscoring the virus’s epidemic potential [1– 3]. Although historically confined to endemic regions in Central and West Africa, MPXV has demonstrated increased transmissibility and geographic spread, raising concerns regarding preparedness for orthopoxvirus re-emergence.

Currently, antiviral options for orthopoxvirus infections are limited. Tecovirimat (ST-246) is the primary approved antiviral agent, targeting the viral VP37 protein, but reports of resistance and limited clinical data necessitate alternative or complementary therapies [4–6]. Brincidofovir and cidofovir exhibit activity against orthopoxviruses but are associated with significant toxicity [7]. Therefore, the identification of novel, orally bioavailable, and well-tolerated antivirals remains a critical unmet need.

Nucleoside and nucleoside analogs have historically played a central role in antiviral drug development. N4-hydroxycytidine (NHC) and its prodrug EIDD-2801 (molnupiravir) exhibit broad-spectrum antiviral activity via lethal mutagenesis, particularly against RNA viruses such as SARS-CoV-2 [8–10]. Previously, we reported potent anti-coronavirus activity of NHC analogs, highlighting structure–activity relationships that enhance antiviral potency. Given the mechanistic versatility of nucleoside analogs, we hypothesized that selected NHC derivatives might also exhibit activity against DNA viruses, including orthopoxviruses [11].

In this study, we investigated the antiviral efficacy of the NHC analog SN_9 against Vaccinia virus (VACV), a well-established surrogate model for MPXV and variola virus. Due to the high genomic homology (>85–86%) and conserved replication machinery among orthopoxviruses, antiviral activity against VACV is highly predictive of efficacy against MPXV [12, 13].

## Methods

### Cells and virus

Vero E6 cells (ATCC CRL-1586) were grown in DMEM containing 10% fetal calf serum (FCS) (Hyclone, USA), 2 mM L-glutamine (Gibco, USA), 100 U/ml penicillin and 100 μg/ml streptomycin (Gibco, USA). A strain of VACV Western Reserve was propagated in Vero E6 cells.

### Viral challenges

Mice BALB/c were anesthetised with zoletil and xylazine by intraperitoneal injections, and the challenge virus was delivered to the nares. The SN_9 was prepared for oral dosing of mice by suspension in an aqueous solution containing 0.75% methylcellulose. SN_9 was delivered in a volume of 100 μL by oral gavage (400 mg/kg). Animals were dosed daily at 24-h intervals for 5 or 10 days beginning on the day of challenge with VV-WR. All animal protocols were approved by the Biomedical Ethics Committee of the Gamaleya Center. Mice were then monitored for survival, signs of disease and weight loss.

## Results

### Inhibition of Vaccinia Virus Replication by SN_9 In Vitro

The antiviral activity of SN_9 was evaluated in cell culture using Vaccinia virus (VACV) strain Lister. Treatment with SN_9 resulted in a pronounced suppression of viral replication, as demonstrated by a significant reduction in virus-induced cytopathic effect (CPE) compared with untreated infected controls. When directly compared with EIDD-2801 (molnupiravir), SN_9 exhibited superior antiviral efficacy, achieving a greater degree of viral inhibition at comparable concentrations (Figure 1).

**Figure 1.**
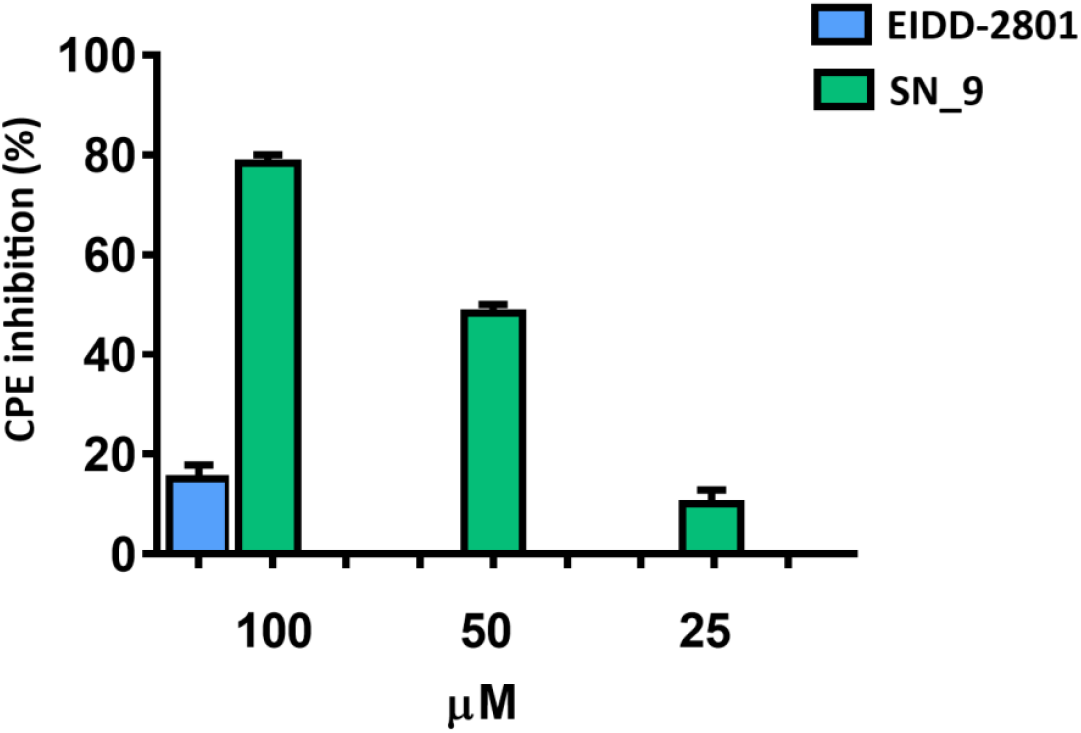
Graphs of inhibition of VACV-induced cytopathic effect (CPE).

**Figure 2.**
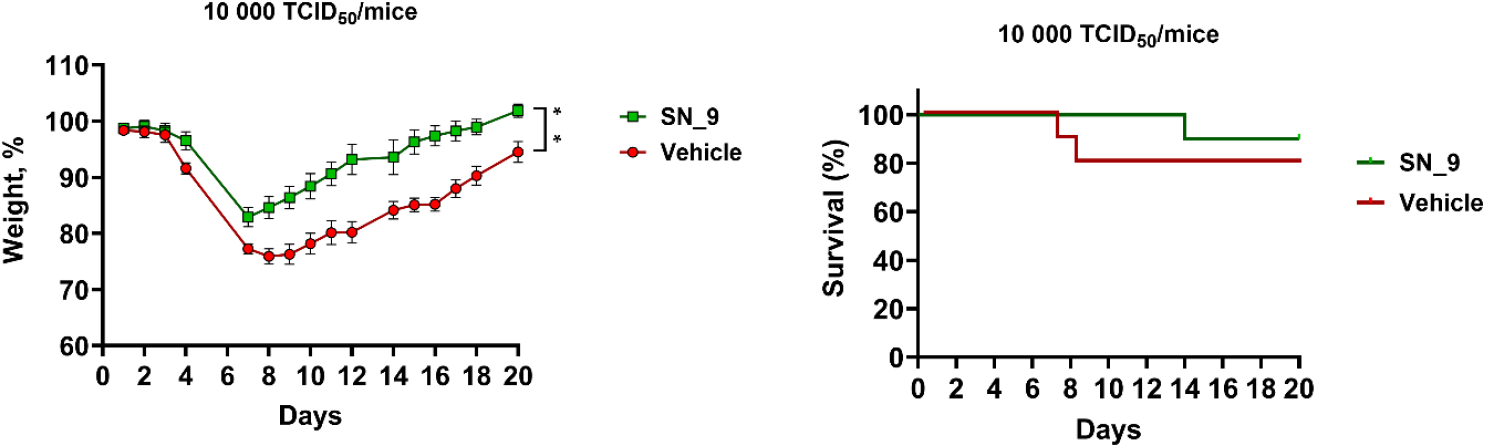
Efficacy of SN_9 in sublethal VV-WR challenge in BALB/c mice. Mice were intranasally challenged (day 0) with the wild-type VV-WR and was treated with SN_9 for 5 consecutive days. Mice were monitored for evidence of disease and were killed if moribund. Survival percentages are shown.

These findings indicate that SN_9 possesses enhanced anti-orthopoxvirus activity relative to its parent compound and suggest improved interaction with viral replication processes. The *in vitro* data provided the rationale for further evaluation of SN_9 in animal models of orthopoxvirus infection. Moreover, structural modifications introduced in SN_9 enhance antiviral activity beyond that of the parent compound molnupiravir, potentially improving interaction with viral replication machinery or increasing intracellular bioavailability.

### Protective Efficacy of SN_9 in a Sublethal Mouse Model of VACV Infection

The *in vivo* antiviral efficacy of SN_9 was first assessed in a sublethal intranasal infection model using BALB/c mice infected with VACV Western Reserve. SN_9 was administered orally twice daily (400 mg/kg, BID) for five consecutive days following infection.

As shown in Figure 4A, animals treated with SN_9 displayed significantly attenuated clinical manifestations of infection. In contrast to infected control mice, which experienced marked body weight loss and delayed recovery, SN_9-treated animals exhibited only transient weight loss followed by rapid restoration of baseline body mass. Importantly, survival in the SN_9 treatment group reached 90%, whereas untreated infected animals demonstrated increased morbidity and mortality.

These results indicate that SN_9 effectively limits disease progression and promotes recovery during orthopoxvirus infection under sublethal conditions.

### SN_9 Improves Survival in a Lethal VACV Challenge Model

To further examine the therapeutic potential of SN_9 under severe disease conditions, a lethal VACV infection model was used. Mice were intranasally infected with a high viral dose and treated orally with SN_9 twice daily (400 mg/kg, BID) for ten days.

As illustrated in Figure 3, SN_9 treatment significantly delayed the onset of clinical disease and reduced overall mortality. While control animals rapidly succumbed to infection, SN_9-treated mice demonstrated a significant increase in survival, with 70% of animals surviving the lethal challenge. Statistical analysis using the log-rank (Mantel–Cox) test confirmed that the observed survival advantage was significant (p < 0.05).

Collectively, these findings demonstrate that SN_9 confers robust protection against orthopoxvirus infection *in vivo*, even under conditions of high viral burden.

**Figure 3.**
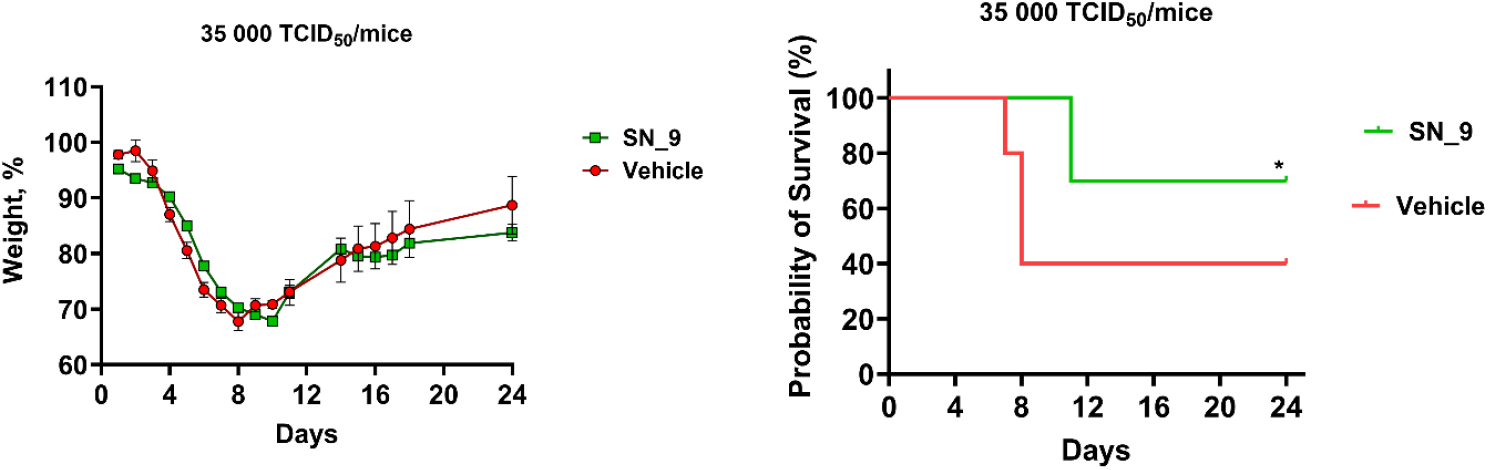
Efficacy of SN_9 in lethal VV-WR challenge in BALB/c mice. Mice were intranasally challenged with the wild-type VV-WR on day 0. Each mouse was treated daily with vehicle or with SN_9 for 10 consecutive days starting at the time of challenge. Survival percentages after challenge and average weights are shown.

## Discussion

The present study demonstrates that the NHC conjugate with phenylpropionic acid (SN_9) exhibits potent antiviral activity against orthopoxviruses both *in vitro* and *in vivo*. Importantly, SN_9 outperformed EIDD-2801 in suppressing VACV replication, suggesting that rational chemical modification of NHC scaffolds can extend antiviral activity beyond RNA viruses.

Orthopoxviruses replicate in the cytoplasm and encode their own DNA-dependent RNA polymerase and DNA replication enzymes, which differ substantially from host counterparts [14,15]. Nucleoside analogs targeting viral polymerases have long been effective against DNA viruses, including herpesviruses and poxviruses [16–18]. Although the precise mechanism of SN_9 against VACV remains to be elucidated, possible modes of action include interference with viral DNA synthesis, transcriptional dysregulation, or induction of mutagenesis.

The use of VACV as a surrogate model is well established, as VACV, MPXV, and variola virus share extensive genomic conservation and similar pathogenic mechanisms [11,19]. Historically, antivirals and vaccines effective against VACV have demonstrated cross-protective efficacy against MPXV [20–22]. Therefore, the observed efficacy of SN_9 against VACV strongly supports its potential utility against Mpox. Notably, SN_9 not only improved survival but also mitigated disease severity, as evidenced by faster recovery of body weight and delayed disease onset. These parameters are critical translational indicators, as weight loss and survival closely correlate with viral load and systemic disease progression in orthopoxvirus infections.

Compared with existing antivirals such as tecovirimat, SN_9 offers a distinct mechanism of action, which could reduce the likelihood of cross-resistance and enable combination therapy [5,6]. Moreover, oral administration and favorable tolerability in animal models further enhance its translational potential.

Given the ongoing circulation of MPXV and the risk of future outbreaks, the development of broad-spectrum orthopoxvirus antivirals remains a global priority. SN_9 represents a promising lead compound for further preclinical development, including pharmacokinetic profiling, toxicity assessment, and direct evaluation against MPXV in relevant animal models.

The strong in vivo efficacy of SN_9 observed in both sublethal and lethal models is particularly relevant in the context of emerging mpox outbreaks. Vaccinia virus is widely accepted as a predictive surrogate for monkeypox virus (MPXV), owing to extensive genomic homology and conservation of essential replication proteins among orthopoxviruses. Therefore, antiviral activity against VACV is strongly indicative of potential efficacy against MPXV. Given that current therapeutic options such as tecovirimat target a single viral protein, the identification of nucleoside analogs with distinct mechanisms of action is of high importance. SN_9 may therefore represent a valuable candidate either as a standalone therapy or as part of combination regimens aimed at reducing resistance development and improving clinical outcomes in Mpox patients.

## Conclusion

The present study demonstrates that the N4-hydroxycytidine analog SN_9 exhibits potent antiviral activity against orthopoxviruses in vitro and in vivo. Oral administration of SN_9 significantly reduced disease severity and mortality in murine models of Vaccinia virus infection. Given the close genetic and functional relatedness of VACV, MPXV, and variola virus, these findings strongly support further development of SN_9 as a potential therapeutic and prophylactic agent for monkeypox. Further mechanistic and translational studies are warranted to advance SN_9 toward clinical evaluations.

## Notes

### Competing Interest Statement

The authors have declared no competing interest.

### Funding

- Ministry of Health of the Russian Federation: 125021101875-3

## References

1. Thornhill JP et al. N Engl J Med. 2022. DOI: 10.1056/NEJMoa2207323

2. Bunge EM et al. PLoS Negl Trop Dis. 2022. DOI: 10.1371/journal.pntd.0010141

3. Reynolds MG, Damon IK. Lancet. 2012. DOI: 10.1016/S0140-6736(11)61762-2

4. Grosenbach DW et al. Nature. 2018. DOI: 10.1038/nature24091

5. Russo AT et al. Antiviral Research. 2021. DOI: 10.1016/j.antiviral.2021.105019

6. Duraffour S et al. Viruses. 2021. DOI: 10.3390/v13081533

7. Parker S et al. Antiviral Research. 2008. DOI: 10.1016/j.antiviral.2008.02.006

8. Sheahan TP et al. Nat Commun. 2020. DOI: 10.1038/s41467-020-18447-9

9. Kabinger F et al. Nat Struct Mol Biol. 2021. DOI: 10.1038/s41594-021-00651-0

10. Malone B, Campbell EA. Nat Struct Mol Biol. 2021. DOI: 10.1038/s41594-021-00657-8

11. Siniavin AE. Antiviral Res. 2024. DOI: 10.1016/j.antiviral.2024.105871

12. Petersen BW et al. MMWR. 2019. DOI: 10.15585/mmwr.mm6810a5

13. Seet BT et al. Nat Rev Microbiol. 2003. DOI: 10.1038/nrmicro754

14. McFadden G. PLoS Pathog. 2005. DOI: 10.1371/journal.ppat.0010025

15. De Clercq E. Clin Microbiol Rev. 2003. DOI: 10.1128/CMR.16.4.569-596.2003

16. Andrei G et al. Antiviral Research. 2006. DOI: 10.1016/j.antiviral.2005.10.008

17. Keith KA et al. Antimicrob Agents Chemother. 2003. DOI: 10.1128/AAC.47.6.1960-1967.2003

18. Carroll DS et al. J Virol. 2012. DOI: 10.1128/JVI.00215-12

19. Hooper JW et al. J Infect Dis. 2004. DOI: 10.1086/422303

20. Earl PL et al. Vaccine. 2004. DOI: 10.1016/j.vaccine.2004.03.035

21. Reynolds MG et al. Emerg Infect Dis. 2010. DOI: 10.3201/eid1606.100010

